# In vitro heterochronic parabiosis identifies pigment epithelium-derived factor as a systemic mediator of rejuvenation by young blood

**DOI:** 10.1101/2024.05.02.592258

**Authors:** Xizhe Wang, Cagdas Tazearslan, Seungsoo Kim, Qinghua Guo, Daniela Contreras, Jiping Yang, Adam D. Hudgins, Yousin Suh

**Affiliations:** Department of Obstetrics and Gynecology, Columbia University Medical Center, New York, NY; Department of Genetics and Development, Columbia University Medical Center, New York, NY; Department of Genetics, Albert Einstein College of Medicine, Bronx, NY

## Abstract

Several decades of heterochronic parabiosis (HCPB) studies have demonstrated the restorative impact of young blood, and deleterious influence of aged blood, on physiological function and homeostasis across tissues, although few of the factors responsible for these observations have been identified. Here we develop an in vitro HCPB system to identify these circulating factors, using replicative lifespan (RLS) of primary human fibroblasts as an endpoint of cellular health. We find that RLS is inversely correlated with serum donor age and sensitive to the presence or absence of specific serum components. Through in vitro HCPB, we identify the secreted protein pigment epithelium-derived factor (PEDF) as a circulating factor that extends RLS of primary human fibroblasts and declines with age in mammals. Systemic administration of PEDF to aged mice reverses age-related functional decline and pathology across several tissues, improving cognitive function and reducing hepatic fibrosis and renal lipid accumulation. Together, our data supports PEDF as a systemic mediator of the effect of young blood on organismal health and homeostasis and establishes our in vitro HCPB system as a valuable screening platform for the identification of candidate circulating factors involved in aging and rejuvenation.

---

Studies of heterochronic parabiosis (HCPB) and heterochronic plasma transfer have demonstrated the rejuvenation of aged tissue by exposure to young blood, and the induction of aging-associated dysfunction in young tissue by exposure to aged blood, across tissues, including the brain, liver, kidney, muscle, heart, bone, spinal cord, pancreas, ovary, vasculature, and skin^1–15^. HCPB studies have also reported the age-dependent influence of the circulation on the extent of cellular senescence across tissues with age^16^, as well as epigenetic clock measures of tissue biological age and lifespan^17^. However, the identification of the specific circulating factors that mediate these phenotypes has remained limited, partly due to the time, effort, and resources necessary to evaluate candidates in animal models. Establishing a high-throughput screening method requires a system, such as in vitro cell culture, which is amenable to scaling, has ease of accessibility, and has a shorter timeframe required for evaluation. Cell culture also provides a powerful tool for studying mechanisms of cellular health.

Recent progress in geroscience indicates great potential for delaying the onset and progression of multiple aging-related diseases by targeting basic aging biology^18–20^. The so-called “geroprotectors”, such as rapamycin^21^, that target conserved mechanisms of aging and extend lifespan and healthspan across taxa^22^, have also been shown to attenuate in vitro aging of human mesenchymal stem cells by enhancing proliferative capacity, self-renewal, and osteogenic potential^23^, as well as to delay certain aspects of cellular senescence, including extension of replicative lifespan (RLS) of human primary fibroblasts^24–28^. Thus, we reasoned that RLS of human primary fibroblast culture could be harnessed as a surrogate endpoint of geroprotection to develop an in vitro HCPB circulating factor screening system.

## RLS of human primary fibroblasts is sensitive to serum age

The effect of sera donor age on fibroblast growth in vitro was first documented in the 1920s, where it was found that young serum was more beneficial for the growth of chicken fibroblasts than aged serum^29^. More recently, it has been shown that exposure to aged serum leads to a decrease in myotube diameter in human skeletal muscle myotube cultures, suggesting that cell culture using serum of different ages can be used as a model to study age-related muscle loss^30,31^. As the initial step in the development of our in vitro HCPB screening system we first determined the impact of serum age on in vitro cellular health and RLS and tested the effects of culturing IMR90 human primary fibroblasts in bovine sera of different ages (fetal (FBS), 1.5 years, 3 years, 6 years, and 10 years of age). We observed that RLS is inversely correlated with the age of serum donor (**Fig. 1a**), with culture in 10-year-old bovine serum, approximately equivalent to that of a 50-year-old human^32^, resulting in a significantly shorter RLS when compared to culture in fetal serum. In mid-late passages, cells cultured in aged serum exhibited increased senescence as indicated by loss of LaminB1 (*LMNB1*) expression (**Fig. 1b**), induction of p16^INK4A^ (*CDKN2A*) expression (**Fig. 1c**), a higher percentage of senescence-associated-β galactosidase (SA-β-gal) positivity (**Fig 1d-e**), and a lower percentage of actively replicating cells as indicated by Ki67 positivity (**Fig. 1f-g**). To gain greater insight into the impact of serum age on in vitro cellular health we performed RNA-seq analysis of early passage IMR90 cultures 24 hours post-serum change into either fetal or aged (10y) serum and found 771 differentially expressed genes (DEGs) (**Fig. 1h and Supplementary Table 1**). Enrichment analysis of Molecular Signatures Database Hallmark gene sets showed that the downregulated DEGs between aged and fetal serum conditions were enriched for genes involved in cell cycle regulation and mitosis (**Fig. 1i**). Conversely, the upregulated DEGs were enriched for processes involved in cell stress, inflammation, and senescence (**Fig. 1j**). Consistent with the qPCR analysis, p16^INK4A^ (*CDKN2A*) expression was also increased after exposure to aged serum, as were many of the genes from the recently defined SenMayo^33^ senescence marker gene set (**Fig. 1k**). Taken together, these results indicate that the differential composition of sera due to age has direct and robust impacts on cellular health and lifespan in vitro.

**Figure 1.**
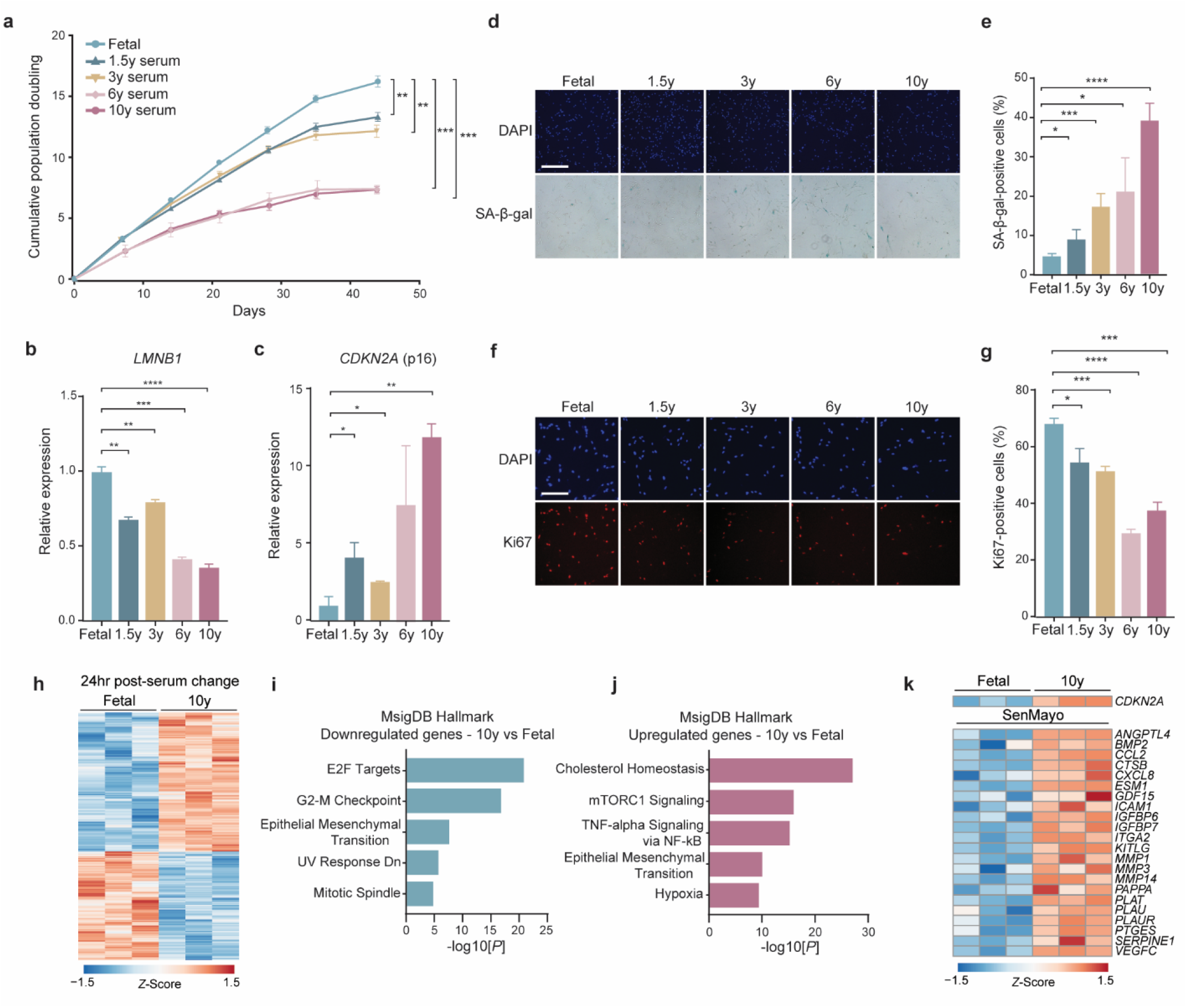
Aged serum reduces replicative lifespan and induces early senescence of human primary fibroblasts. **a**, Replicative lifespan curves of human primary fibroblast, IMR90, under different bovine serum conditions from different ages (fetal, 1.5y, 3y, 6y, and 10y). **b**, qPCR analysis of *LMNB1* in mid-late passage IMR90 cultured under different serum age conditions, relative to fetal serum. **c**, qPCR analysis of *CDKN2A* in mid-late passage IMR90 cultured under different serum age conditions, relative to fetal serum. **d-e**, Representative images (**d**) and quantification (**e**) of senescence-associated β-galactosidase (SA-β-Gal) staining of IMR90 at mid-late passage from different serum age conditions. **f-g,** Representative images (**f**) and quantification (**g**) of *Ki67* staining of IMR90 at mid-late passage from different serum age conditions. **h-k**, Significant DEGs (FDR<0.05) after RNA-seq analysis of IMR90 24hr post-serum change from fetal to 10-year-old serum (**h**), associated pathways (**i-j**), and expression of *CDKN2A* and SenMayo senescence genes among significant DEGs (**k**). Statistical analysis was performed using two-tailed unpaired *t*-tests (**a-c**,**e** and **g**), Wald tests (**h** and **k**), and Fisher’s exact test (**i-j**). Data represented as mean ±SEM from three technical replicates, \**p*<0.05, \*\**p*<0.01, \*\*\**p*<0.001, \*\*\*\**p*<0.0001. Scale bars, 500 µm (**d**) and 200 µm (**f**).

## Young serum counters negative impact of aged serum on RLS

Next, we mimicked the systemic environment of in vivo HCPB, and cultured IMR90 in half young (5% fetal) and half aged (5% 10y) bovine serum in heterochronic cultures. We observed that the RLS of IMR90 in heterochronic serum culture is significantly extended when compared to the aged serum condition (**Extended Data Fig. 1a**). In mid-late passages, cells cultured in the heterochronic condition had reduced loss of LaminB1 (*LMNB1*) expression as compared to the aged serum cultures (**Extended Data Fig. 1b**), non-significant induction of p16^INK4A^ (*CDKN2A*) expression as compared to the young serum condition (**Extended Data Fig. 1c**), reduced SA-β-gal positivity in comparison to aged serum cultures (**Extended Data Fig 1d-e**), and a higher percentage of Ki67 positivity as compared to the aged serum condition (**Extended Data Fig. 1f-g**).

However, the chimeric environment achieved by in vivo HCPB involves aged and young blood from postnatal animals, not fetal stage blood. Thus, to more closely mimic in vivo HCPB, we next utilized young adult (1.5 years old) serum as the young serum in the heterochronic cultures. Consistent with the previous results, we observed that the RLS of IMR90 in heterochronic serum culture is significantly extended when compared to the aged serum condition (**Fig. 2a**). In mid-late passages, cells cultured in the heterochronic condition had reduced loss of LaminB1 (*LMNB1*) expression as compared to the aged serum condition (**Fig. 2b**), no induction of p16^INK4A^ (*CDKN2A*) expression as compared to young serum cultures (**Fig. 2c**), attenuated SA-β-gal positivity in comparison to the aged serum condition (**Fig. 2d-e**), and no difference in the percentage of Ki67 positivity as compared to young serum cultures (**Fig. 2f-g**). To translate this system into a human-relevant context, we next created isochronic and heterochronic cultures using sex-specific pooled human serum from young (23-29 years) and aged (58-65 years) donors, both male and female. Consistent with the findings from bovine serum, we observed that culturing IMR90 in aged human serum resulted in reduced RLS, and that this reduction was rescued in the heterochronic condition (**Fig. 2h-i**). In the male heterochronic cultures RLS was indistinguishable from that of young serum cultures (**Fig. 2h**), while in the female heterochronic cultures there was only a partial rescue of RLS, in comparison to the aged serum condition (**Fig. 2i**). These data indicate that RLS in our in vitro HCPB system is responsive to the relative age and composition of the serum used in the culture medium, and that this system can be used as a model of a conserved cellular response to the systemic milieu.

**Figure 2.**
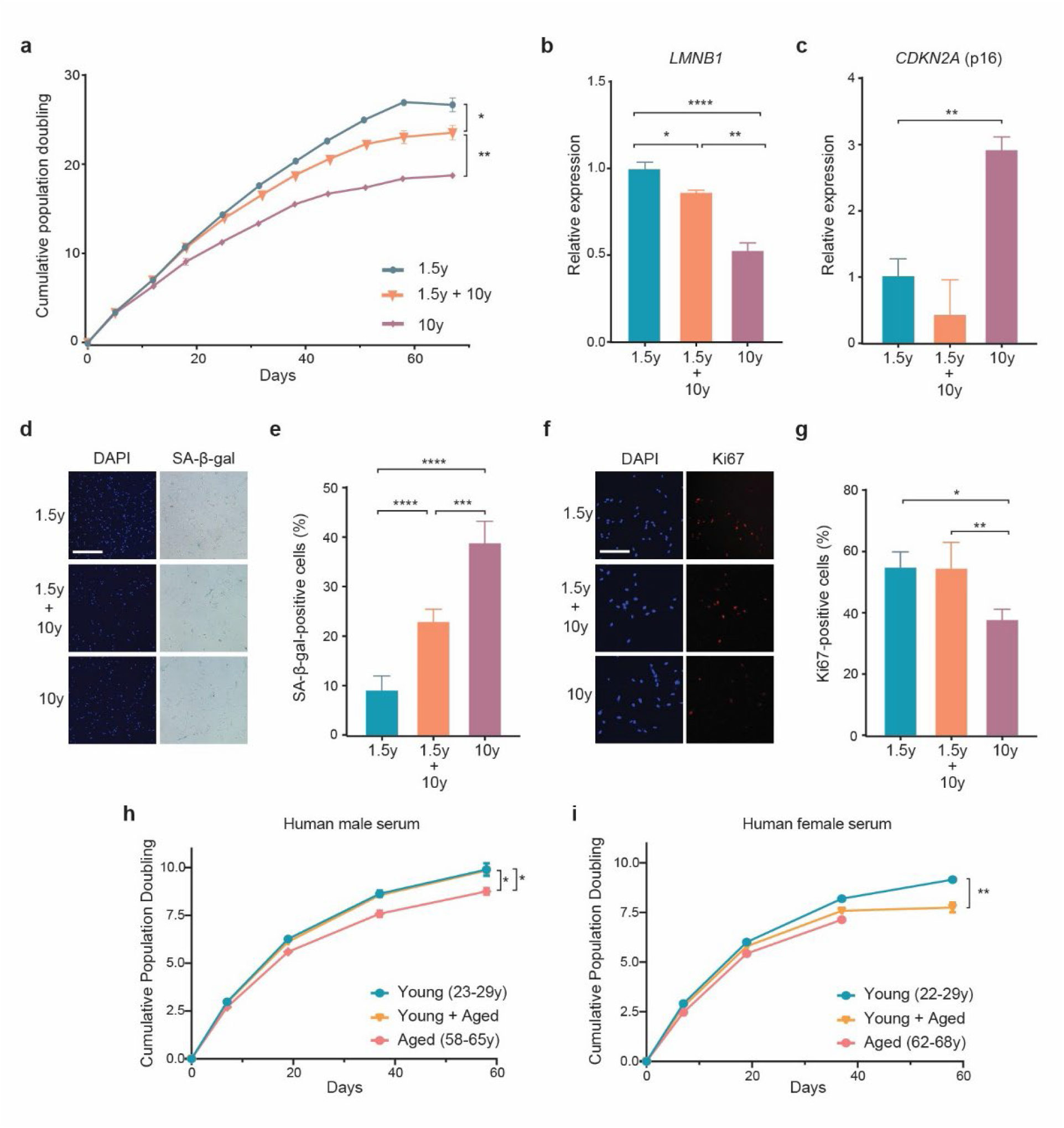
Young serum counteracts deleterious impact of aged serum in heterochronic culture system. **a,** Replicative lifespan curves of IMR90 cultured in young (1.5y), aged (10y), and heterochronic (50% young/50% aged) bovine serum. **b**, qPCR analysis of *LMNB1* in mid-late passage IMR90 cultured under different serum conditions, normalized to young serum condition (1.5 years). **c**, qPCR analysis of *CDKN2A* in mid-late passage IMR90 cultured under different serum conditions, normalized to young serum condition (1.5 years). **d-e**, Representative images (**d**) and quantification (**e**) of senescence-associated β-galactosidase (SA-β-Gal) staining of IMR90 at mid-late passage from different serum age conditions. **f-g,** Representative images (**f**) and quantification (**g**) of *Ki67* staining of IMR90 at mid-late passage from different serum age conditions. **h-i**, Replicative lifespan curves of IMR90 cultured in young (22-29y), aged (58-68y), and heterochronic (50% young/50% aged) male (**h**) and female (**i**) human serum. Statistical analysis was performed using two-tailed unpaired *t*-tests (**a-c**,**e** and **g-i**). Data represented as mean ±SEM from three technical replicates, \**p*<0.05, \*\**p*<0.01, \*\*\**p*<0.001, \*\*\*\**p*<0.0001. Scale bars, 500 µm (**d**) and 200 µm (**f**).

## Serum protein factors have the strongest impact on RLS

To maximize the utility of our in vitro HCPB system we sought to identify which class of molecules in serum had the greatest impact on RLS, so that they could be prioritized for screening in supplementation or depletion experiments. Fetal bovine serum (FBS) contains a rich assortment of hormones, growth factors, amino acids, proteins, vitamins, inorganic salts, trace elements, carbohydrates, lipids, and other metabolites that are essential for cellular metabolism and growth^34^. To test the importance of these different serum components to RLS of human primary fibroblast cultures, we utilized different processed forms of FBS that had specific fractions of factors selectively removed. This included dialysis to remove metabolites, amino acids, and micronutrients; exosome depletion to eliminate miRNAs and other exosome cargo; charcoal stripping to remove non-polar molecules including lipids and hormones; and heat inactivation to partially (76 °C) or irreversibly (95 °C) denature proteins.

We supplemented these processed FBS samples (5% concentration of each sample) to IMR90 cell culture media containing 5% standard FBS (10% total serum concentration) and measured impact on RLS. We found that, except for the dialyzed serum, all other processed FBS cultures had significantly reduced RLS, in comparison to the standard FBS condition (**Fig. 3a**). The heat-inactivated FBS cultures had the most pronounced reduction in RLS, both in the 76 °C treated cultures and in the 95 °C treated cultures, which only reached a fraction of the RLS observed in the other serum conditions (**Fig. 3a**). In line with these observations, at mid-late passage the heat-inactivated FBS groups also showed the strongest markers of senescence onset, as indicated by the greatest loss of LaminB1 (*LMNB1*) expression (**Fig. 3b**), the highest induction of p16^INK4A^ (*CDKN2A*) expression (**Fig. 3c**), the highest percentage of SA-β-gal positivity (**Fig 3d-e**), and the lowest percentage of actively replicating cells as indicated by Ki67 positivity (**Fig. 3f-g**). Collectively, these results emphasize the importance of serum protein factors in the modulation of phenotypes of cellular senescence and RLS of human primary fibroblast cultures and prioritized the protein fraction for further analyses.

**Figure 3.**
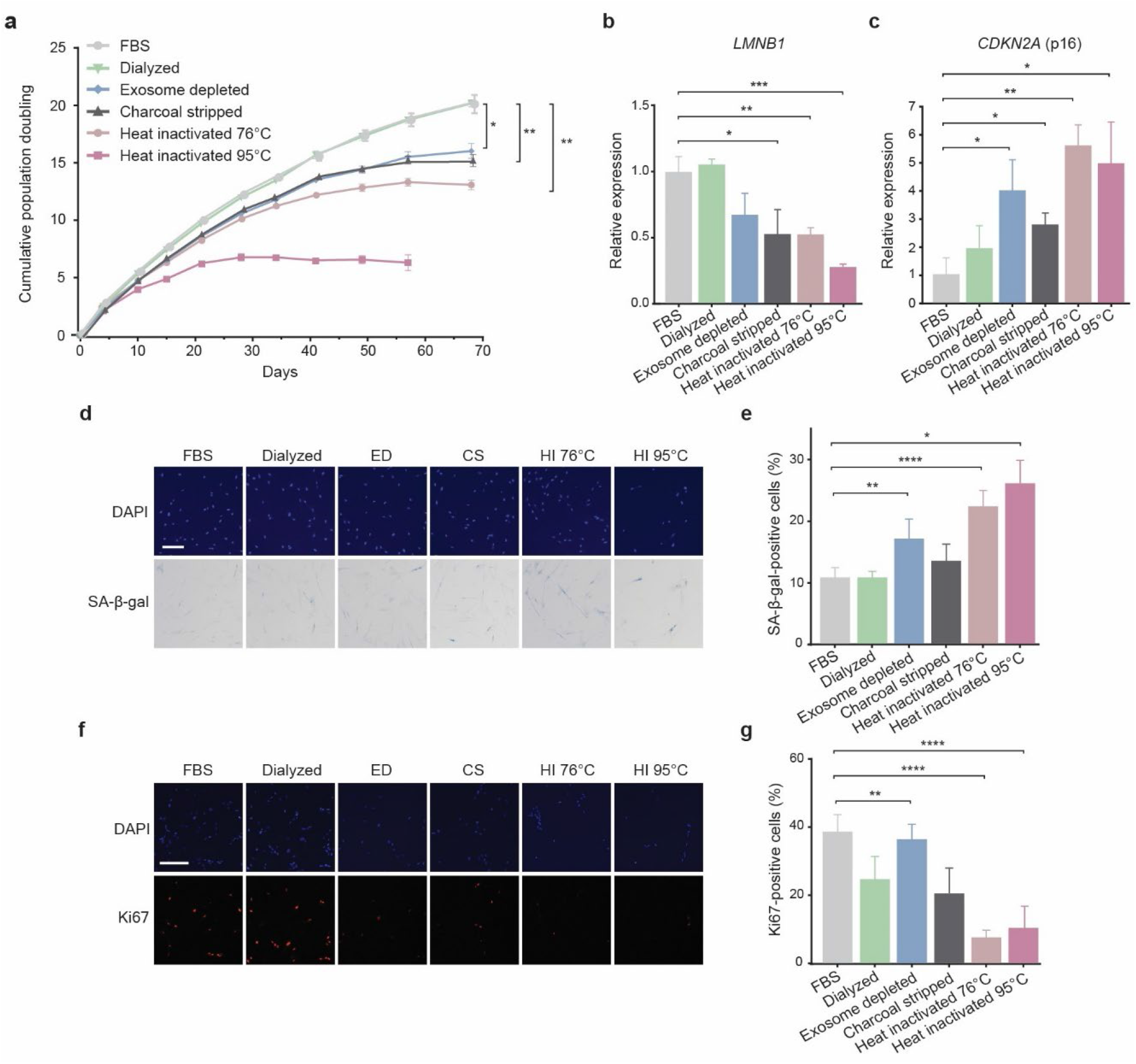
Serum proteins have the strongest impact on replicative lifespan. **a,** Replicative lifespan curves of IMR90 cultured in fetal serum subjected to different fraction depletion/inactivation manipulations. **b,** qPCR analysis of *LMNB1* in mid-late passage IMR90 cultured in different serum conditions, normalized to unmanipulated fetal serum. **c**, qPCR analysis of *CDKN2A* in mid-late passage IMR90 cultured in different serum conditions, normalized to unmanipulated fetal serum. **d-e**, Representative images (**d**) and quantification (**e**) of senescence-associated β-galactosidase (SA-β-Gal) staining of IMR90 at mid-late passage in different serum conditions. **f-g,** Representative images (**f**) and quantification (**g**) of *Ki67* staining of IMR90 at mid-late passage in different serum age conditions. Statistical analysis was performed using two-tailed unpaired *t*-tests (**a-c**,**e** and **g**). Data represented as mean ±SEM from three technical replicates, \**p*<0.05, \*\**p*<0.01, \*\*\**p*<0.001, \*\*\*\**p*<0.0001. Scale bars, 200 µm (**d** and **f**). ED=exosome depleted; CS=charcoal stripped; HI=heat inactivated

## Proteomic analysis identifies candidate circulating factors

To investigate the changes in circulating protein factors during aging as a means to identify and prioritize candidate factors for testing in our in vitro screening system, we utilized data-independent acquisition (DIA) quantitative mass spectrometry to analyze bovine sera of different ages (fetal, 1.5 years old, 3 years old, 6 years old, and 10 years old). Principal component analysis (PCA) of the DIA mass spectrometry results revealed that fetal bovine serum is compositionally distinct as compared to the other age groups (**Extended Data Fig. 2a**), which collectively showed a distribution that corresponded to sample age. Based on these results, we decided to exclude fetal bovine serum from further analysis due to its distinct composition and restricted our analysis to the samples taken over the life course of postnatal animals. Examining the 647 proteins that were identified across all samples, we found that, overall, the levels of most proteins in bovine serum tend to decrease with age (**Fig. 4a**). To further investigate the shifts in abundance across age, protein trajectories were estimated using LOESS regression (**Fig. 4b**), and the resulting trajectories were subjected to unsupervised hierarchical clustering **(Fig. 4c**), which identified 10 clusters of protein trajectories (**Fig. 4d, Supplementary Table 2**), most of which were enriched for biological pathway terms, indicating that particular aging trajectories were associated with specific biological processes (**Fig. 4d**). Notably, the majority of proteins in bovine serum did not exhibit a linear change in abundance with age, consistent with similar trends observed in other studies examining circulating protein levels^35–38^, including data from humans and mice, which indicates conservation of this phenomenon. This nonlinear pattern indicated that many serum proteins do not undergo a gradual and predictable alteration in protein abundance as individuals age, although the proteins in the cluster that contained the largest fraction of proteins, cluster C4 (n=156) did appear to undergo a relatively monotonic decrease in abundance with age (**Fig. 4d**). We then focused on proteins that exhibited simple linear changes in abundance with age and applied a standard linear model, identifying 81 proteins whose abundance significantly changed with age (**Supplementary Table 3**).

**Figure 4.**
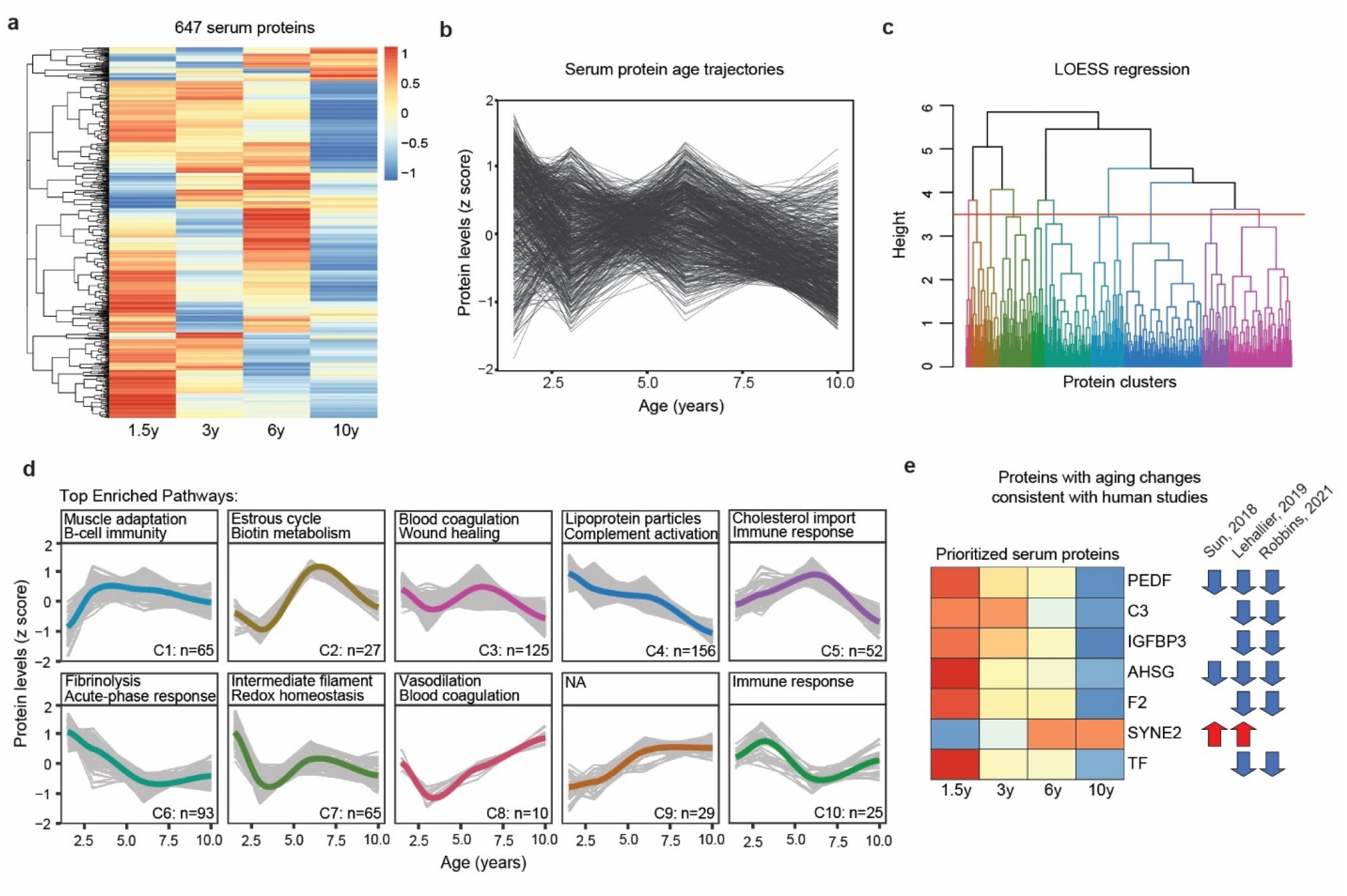
Quantitative bovine serum proteomics identifies conserved protein aging trajectories. **a**, Quantitative DIA mass spectrometry detected 647 bovine serum proteins across ages. Heatmap displays average abundance across technical replicates for each sample age. **b**, Serum protein aging trajectories. Protein levels were z scored and trajectories were estimated by LOESS. **c**, Hierarchical clustering dendrogram of protein trajectories estimated by LOESS regression. **d**, Protein trajectories for the 10 identified clusters. Clusters are grouped by similarity of trajectory, with the thick colored lines representing the average trajectory across proteins in that cluster. The number of proteins in each cluster and the top enriched Gene ontology: biological process pathways determined by Fisher’s exact test are indicated. **e**, Conserved serum proteins with significant differential abundance with age in both our bovine serum proteomic analysis and across 3 public large-scale human plasma/serum studies. Proteins ranked by statistical significance as determined by a linear model from top to bottom. Arrow direction/color indicates correspondence of abundance change between bovine and human. Blue/down = decreased with age, red/up = increased with age.

To further prioritize these candidate circulating protein factors for testing in our heterochronic culture system, we next searched for cross-species conservation of abundance change with age between our bovine serum proteomic data and data from three large-scale human cohort plasma proteomics studies^38–40^. Using the criteria of consistent change in direction in at least two of the three human datasets, we identified 7 circulating proteins with consistent and conserved shifts in abundance with age (**Fig. 4e**). Six of these 7 proteins showed significantly decreased abundance in the circulation with age (PEDF, C3, IGFBP3, AHSG, F2 and TF), and one showed significant increase in abundance with age (SYNE2) (**Fig. 4e**). From these prioritized candidates, our top candidate pigment epithelium-derived factor (PEDF) had the most significant decrease in abundance with age in our bovine data, and furthermore was one of only two factors that was found as significantly changed with age across all three human datasets. These data indicate that complex, non-linear shifts in circulating protein abundance is a conserved phenomenon in mammals, and that conserved aging signatures with potential impacts on systemic influences on organismal health can be identified.

## PEDF extends RLS of cells in aged serum

We next wanted to investigate if individual factors identified from our proteomic analysis could mediate the extension of human primary fibroblast RLS that we observed after exposure of cells to young serum. Our top candidate pigment epithelium-derived factor (PEDF) is a conserved multifunctional secreted protein encoded by the *SERPINF1* gene that is widely expressed across tissues in vertebrates (**Extended Data Figure 3a-b**)^41^. Studies have shown that PEDF expression decreases with cellular senescence in vitro^42^, and is lost during human skin aging in vivo^43^. We chose to test the effects of PEDF on the in vitro cellular lifespan of primary human fibroblasts by supplementing to IMR90 cultured in aged human serum, as we had already observed that aged human serum shortens IMR90 RLS (**Fig. 2h-i**). To determine the appropriate dose for PEDF supplementation, we first measured the difference in PEDF serum concentration between young (23-29y) and aged (58-85) male human serum samples by ELISA (**Fig. 5a**). Although the difference in mean PEDF abundance between young and aged serum (4 µg ml^-1^) did not reach statistical significance, likely due to sample size, we chose this concentration as the dosage for PEDF supplementation to aged human serum cultures. We found that PEDF supplementation significantly increased the relative cell viability of IMR90 cultured in the serum of our oldest human sample (85 years, male) (**Fig. 5b**), as well as significantly extending RLS (**Fig. 5c**), in comparison to the aged serum alone, which exhibited an RLS far below that of the young serum (29y, male) cultures.

**Figure 5.**
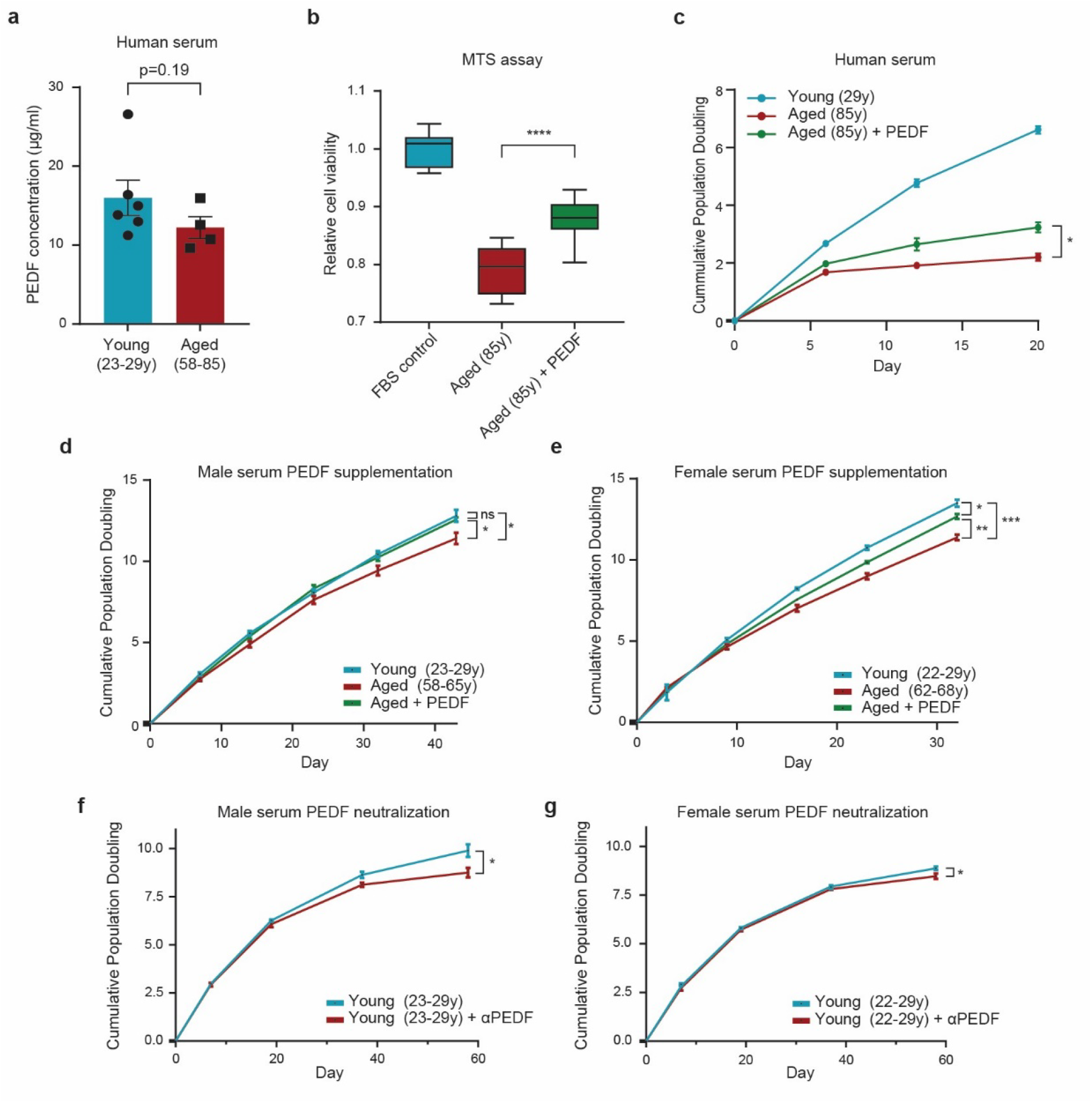
PEDF increases cell viability and replicative lifespan of human primary fibroblasts exposed to aged serum. **a**, ELISA of PEDF in young (23-29y, n=6) and aged (58-85y, n=4) human serum used for replicative lifespan assays. **b-c**, Relative cell viability (**b**) and replicative lifespan curves (**c**) of IMR90 cultured in very aged (85y, male) human serum with and without PEDF supplementation (4 µg ml^-1^), as compared to young human serum (29y, male). **d-e**, Replicative lifespan curves of IMR90, cultured in young (23-29y), aged (58-68y), and aged + PEDF (4 µg ml^-1^) male (**d**) and female (**e**) human serum. **f-g**, Replicative lifespan curves of IMR90 cultured in young (22-29y) and young + PEDF neutralizing antibody male (**f**) and female (**g**) human serum. Statistical analysis was performed using two-tailed unpaired *t*-tests (**a-e**). Data represented as mean ±SEM from three technical replicates, \**p*<0.05, \*\**p*<0.01, \*\*\**p*<0.001, \*\*\*\**p*<0.0001, ns=non-significant.

Since the extreme difference in RLS we observed between the young (29y, male) and aged (85y, male) human serum cultures could be due to individual sample-specific effects on cellular health, we again utilized sex-specific pooled human serum from young (23-29y) and aged (58-65) individuals to further investigate the impact of PEDF on RLS of IMR90. We again observed a significant extension of RLS in the aged serum cultures, both male (**Fig. 5d**) and female (**Fig. 5e**), when supplemented with exogenous PEDF, even though the relative difference in RLS between the pooled young and aged cultures was far less dramatic than what we had observed at the individual serum level (**Fig. 5c**). RNA-seq analysis of early passage IMR90 cultures 24 hours post-serum change into young, aged, and aged supplemented with PEDF human serum cultures found 575 DEGs between the male young and aged serum conditions (**Extended Data Fig. 4a and Supplementary Table 4**), and 683 DEGs between the female young and aged serum conditions (**Extended Data Fig. 4d and Supplementary Table 5**), with no significant DEGs observed in the aged cultures of either sex after PEDF supplementation. Enrichment analysis of Molecular Signatures Database Hallmark gene sets showed that downregulated DEGs between aged and young serum conditions for both sexes were enriched for genes involved in cell cycle regulation and mTORC1 signaling (**Extended Data Fig. 4b-c**). Conversely, upregulated DEGs for both sexes were enriched for processes involved in cell stress, inflammation, and epithelial mesenchymal transition (**Extended Data Fig. 4e-f**). To determine if PEDF is necessary for the observed beneficial effect of young serum on RLS of IMR90, we employed a neutralizing antibody to neutralize PEDF in the young serum and observed a significant reduction in the RLS of the young serum cultures, both male (**Fig. 5f**) and female (**Fig. 5g**). Although acute PEDF treatment of IMR90 cultured in aged human serum did not significantly impact short-term transcriptional outcomes, our findings highlight an important role for PEDF in the RLS of human primary fibroblasts. Moreover, our results demonstrate that PEDF is both necessary and sufficient for the protective effect of young serum on RLS.

## PEDF improves cognition in aged mice

To explore the potential in vivo youth-promoting effects of PEDF, aged (20 months old) male mice received either saline or recombinant PEDF (4 µg day^-1^) through subcutaneous (s.c.) mini-osmotic pumps for a duration of 6 weeks (**Fig. 6a**) and cognitive function and transcriptional changes were assessed. The s.c. mini-osmotic pump route of administration was chosen because of its ability to achieve continuous PEDF infusion, while avoiding drastic spikes in PEDF level, and because of the reduced stress compared to repeated injections, as a single pump implantation surgery was performed^44^. We first tested for the effects of PEDF treatment on cognitive function, as it is one of the most robust phenotypes impacted by HCPB and heterochronic plasma transfer studies^4–6,12,45–47^, and the neurotrophic and neuroprotective functions of PEDF have been well documented^48–50^. Hippocampal-dependent memory was assessed using the novel object recognition (NOR) test after 5 weeks of PEDF treatment and a significant preference for the novel object was observed in the PEDF-treated mice (**Fig. 6b**), with no such preference detected in the saline-treated control animals. To investigate the molecular changes induced by systemic PEDF treatment, we performed RNA-seq analysis of the hippocampus of aged mice after treatment (**Fig. 6c and Supplementary Table 6**). Gene ontology enrichment analysis indicated that DEGs were enriched for processes involving synaptic transmission and cellular responses to factors implicated in brain aging, including iron ions^51^ and amyloid-beta^52^ (**Fig. 6d**). PEDF treatment also increased expression of the metalloproteinase inhibitor *Timp2*, which has been shown to improve cognitive function in the context of heterochronic plasma transfer^12^, the neurotrophic peptide insulin-like growth factor 2, *Igf2*, which has important roles in memory formation^53,54^, and prostaglandin D2 synthase, *Ptgds*, a regulator of circadian rhythm^55^, and oligodendrogenesis and myelination^56^, a critical nervous system process that PEDF has also been shown to influence^57^.

**Figure 6.**
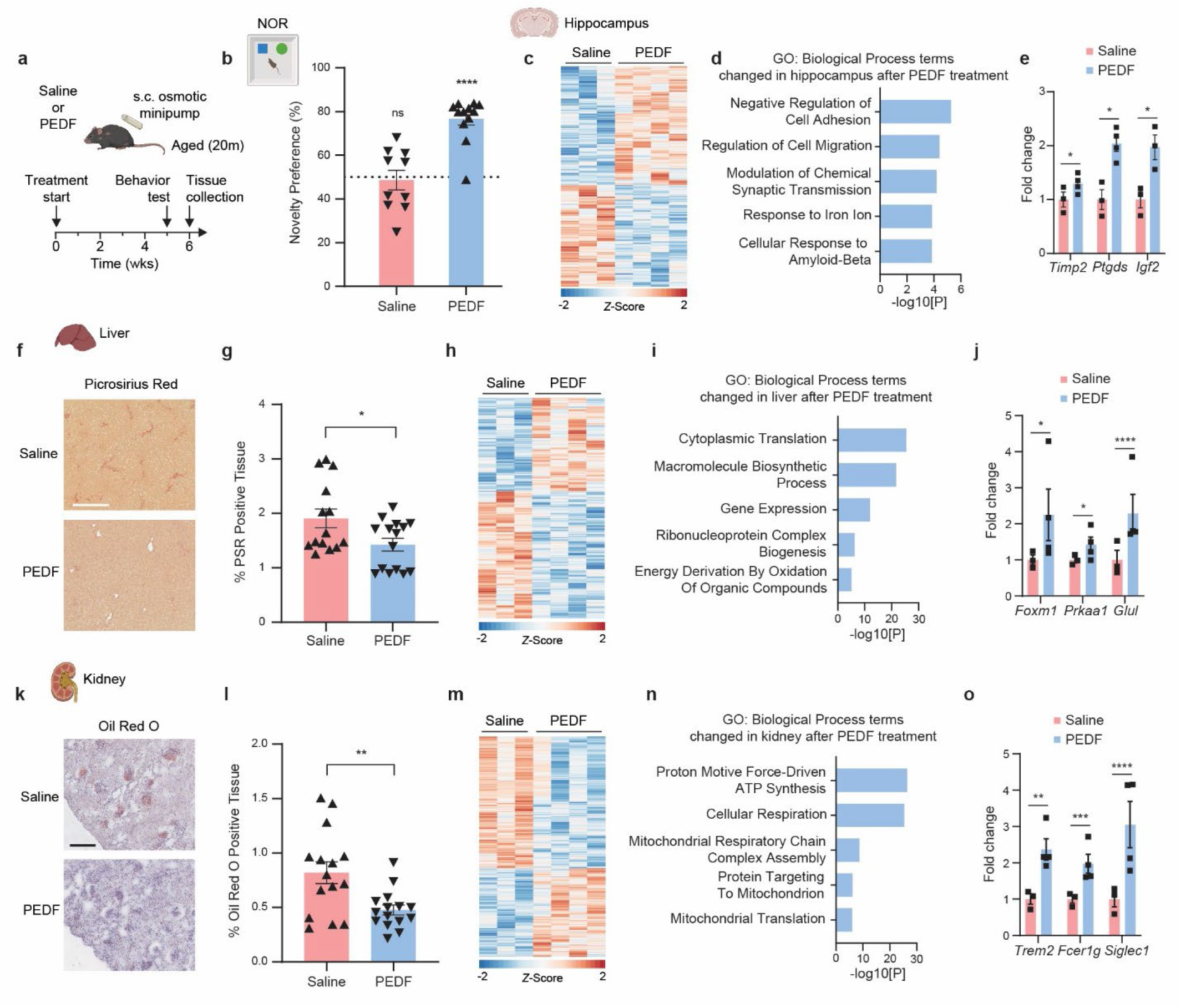
Systemic PEDF improves age-related phenotypes across several tissues in mice. **a,** Schematic of treatment administration timeline to aged (20 months) male mice. s.c., subcutaneous. **b**, Object recognition memory was assessed by Novel Object Recognition (NOR) test as the preference index for the novel object (*n* = 10 (saline), and 12 (PEDF) mice). **c-e**, Significant DEGs (*p*<0.05) after RNA-seq analysis (**c**), associated GO:BP terms (**d**), and the fold change in expression (DEseq2 normalized counts) of example genes among significant DEGs (**e**) in hippocampus from aged mice (n=3 (saline) and 4 (PEDF) mice). **f-g**, Representative images (**f**) and quantification (**g**) of Picrosirius Red staining in liver of aged mice (n=14 (saline), n=15 (PEDF) mice). **h-j**, Significant DEGs (*p*<0.05) after RNA-seq analysis (**h**), associated GO:BP terms (**i**), and the fold change in expression (DEseq2 normalized counts) of example genes among significant DEGs (**j**) in liver from aged mice (n=3 (saline) and 4 (PEDF) mice). **k-l**, Representative images (**k**) and quantification (**l**) of Oil Red O staining in kidney of aged mice (n=15 (saline), n=15 (PEDF) mice). **m-o**, Significant DEGs (*p*<0.05) after RNA-seq analysis (**m**), associated GO:BP terms (**n**), and the fold change in expression (DEseq2 normalized counts) of example genes among significant DEGs (**o**) in kidney from aged mice (n=3 (saline) and 4 (PEDF) mice). Statistical analysis was performed using two-tailed one-sample *t*-test compared to theoretical mean of 0 (**b**), Wald tests (**e**,**j**, and **o**), Fisher’s exact test (**d**,**i** and **n**), and two-tailed unpaired *t*-tests (**g** and **i**). Data represented as mean ±SEM, \**p*<0.05, \*\**p*<0.01, \*\*\**p*<0.001, \*\*\*\**p*<0.0001, ns=non-significant. Scale bars, 500 µm (**g** and **i**).

## PEDF reverses age-related liver and kidney pathology

Studies of HCPB have shown that young blood has a restorative effect on phenotypes of mouse hepatic^1,58^ and renal aging^10^. Based on these findings, we next examined the effects of PEDF treatment on the liver and kidney. Histological analysis revealed a significant reduction in liver fibrosis in aged PEDF-treated mice, as compared to saline-treated controls (**Fig. 6f-g**). No significant difference was observed in terms of hepatic lipid accumulation in aged PEDF-treated mice (**Extended Data Fig. 5a**), suggesting that the positive effects of PEDF on the resolution of liver fibrosis may be independent from processes impacting hepatic steatosis or that the weak steatosis observed in naturally aged mice fed a standard diet^59^ is not responsive to PEDF treatment. Investigation of the effect of PEDF treatment on molecular changes in the aged mouse liver by RNA-seq (**Fig. 6h and Supplementary Table 7**) showed that DEGs were largely enriched for processes involving protein translation and macromolecule anabolism, which were largely downregulated in PEDF-treated livers, as compared to saline-treated controls (**Fig. 6i**).

We also observed that PEDF treatment increased the expression of the transcription factor *Foxm1*, which is a regulator of hepatic regeneration and recovery from injury and disease^60^, *Prkaa1*, a subunit of the classic regulator of metabolic homeostasis AMPK that has been shown to be protective against diet-induced obesity and liver pathology^61^, and the gene encoding glutamine synthetase, *Glul*, which is important for systemic homeostasis and hepatic regulation of brain function^62^ (**Fig. 6j**). These results indicate that the transcriptional signature of reduced protein translation we observed in the PEDF-treated aged liver could be tied to activation of AMPK, which is a well-characterized negative regulator of anabolic pathways^63^. Furthermore, the improved cognitive function observed in PEDF-treated mice (**Fig. 6b**) might be due in part to increased circulating glutamine levels resulting from increased *Glul* expression^64^.

Histological analysis of the kidneys of PEDF- and saline-treated aged mice showed a significant reduction in age-related lipid accumulation in the PEDF-treated group (**Fig. 6k-l**), suggesting that PEDF may play a protective role in countering age-related renal lipotoxicity, which can contribute to the development of tubulointerstitial fibrosis^65^. However, histological analysis of the fibrosis burden of aged kidneys did not identify any significant difference between saline and PEDF treatment (**Extended Data Fig. 5b**). To gain molecular insight into the effects of PEDF treatment on mouse renal aging we performed RNA-seq analysis (**Fig. 6m and Supplementary Table 8**) and found that DEGs between the saline- and PEDF-treated aged kidneys were enriched for gene ontology terms related to ATP synthesis and mitochondrial function (**Fig. 6n**), aspects of cellular metabolism that become dysfunctional in the aged kidney and contribute to excessive generation of reactive oxygen species and inflammation^66^. We observed that PEDF treatment resulted in the increased expression of many genes involved in macrophage biology in the aged kidney (**Supplementary Table 8**). In particular, the expression of genes critical to the regulation of phagocytosis were increased, including *Trem2*, *Fcer1g*, and *Siglec1* (**Fig. 6o**), all of which are involved in the macrophage-mediated scavenging of lipids^67,68^. Considered together, the transcriptional signature of altered mitochondrial function we observed from PEDF treatment (**Fig. 6n**) might be intimately linked with the activation of phagocytic pathways in macrophages, thus resulting in the reduced lipid deposition we observed in the PEDF-treated kidney (**Fig. 6k-l**), as mitochondrial metabolism is a robust regulator of macrophage polarization and function^69,70^.

## Discussion

Overall, our results demonstrate the age-dependent effect of serum on cellular senescence and replicative lifespan in vitro and establish the in vitro HCPB system as a method that can be utilized to screen and investigate circulating factors responsible for the youth- and aging-promoting effects of blood observed in HCPB studies. Our findings highlight the importance of protein factors in mediating the effects of serum age on cellular health and provide further support for the conservation of dynamic serum protein compositional changes with age. In addition, our aging bovine serum quantitative proteomics data should also provide a useful resource for the field. In this study we focused our analyses on the serum protein fraction because it had the strongest impact on RLS when removed, but we also observed that the removal of lipids and hormones and the depletion of exosomes from the serum also significantly impacted RLS of IMR90 cultures. Consistent with our findings, a recent study demonstrated that treatment of aged mice with small extracellular vesicles from the plasma of young mice reversed age-related functional declines in multiple tissues and extended lifespan, which was attributed to vesicle-mediated transfer of miRNA cargoes^71^. These observations provide evidence of the importance of the components of non-protein fractions to the impact of the systemic milieu on organismal homeostasis, components which can be methodically and robustly tested with our in vitro HCPB system.

We identified and characterized PEDF as a conserved circulating geroprotective factor, both in vitro and in vivo, demonstrating that PEDF mediates, in part, the protective effect of young serum on RLS of human primary fibroblasts, and that systemic PEDF administration to aged mice has rejuvenating effects on multiple tissues. Studies of animal models of injury and disease have implicated PEDF in the maintenance of physiological homeostasis through its demonstrated neurotrophic, anti-angiogenic, anti-fibrotic, immunomodulatory, anti-inflammatory, tumoricidal, and stem cell-supporting functions^72–84^. Additionally, PEDF has also been identified as significantly increasing in abundance in plasma following exercise in mice^85^, suggesting that PEDF may play a role in mediating the beneficial effects of exercise on organismal health^86^.

However, there have been no previous studies of the direct effects of PEDF treatment in the context of normal aging. Our results demonstrate that PEDF supplementation induces protective transcriptional signatures in the hippocampus, liver, and kidney of aged mice, improving cognitive function and hepatic and renal integrity, indicating that maintaining or supplementing PEDF levels in aged humans may have therapeutic implications for age-related conditions and chronic diseases of these tissues. Further investigation is needed to understand the mechanisms underlying the impact of PEDF on aged tissue, and insights into the pathways through which PEDF might be promoting systemic homeostasis will be valuable to the development of any future therapies.

## Acknowledgements

We thank members of the Suh lab for their feedback and support. This work was supported by the National Institute on Aging (AG057433). Figure 6 and Extended Data Figure 5 contain elements created with BioRender.com.

## Author contributions

Y.S., A.D.H., X.W., and C.T. conceived and designed the experiments; X.W., C.T., and A.D.H. collected and analyzed data. Q.G., D.C., and J.Y. assisted in data collection. S.K. assisted in data analysis. X.W., A.D.H., and Y.S. wrote the manuscript. A.D.H. and Y.S. supervised all aspects of this project.

## Competing interests

The authors declare no competing interests.

---

**Extended Data Figure 1.**
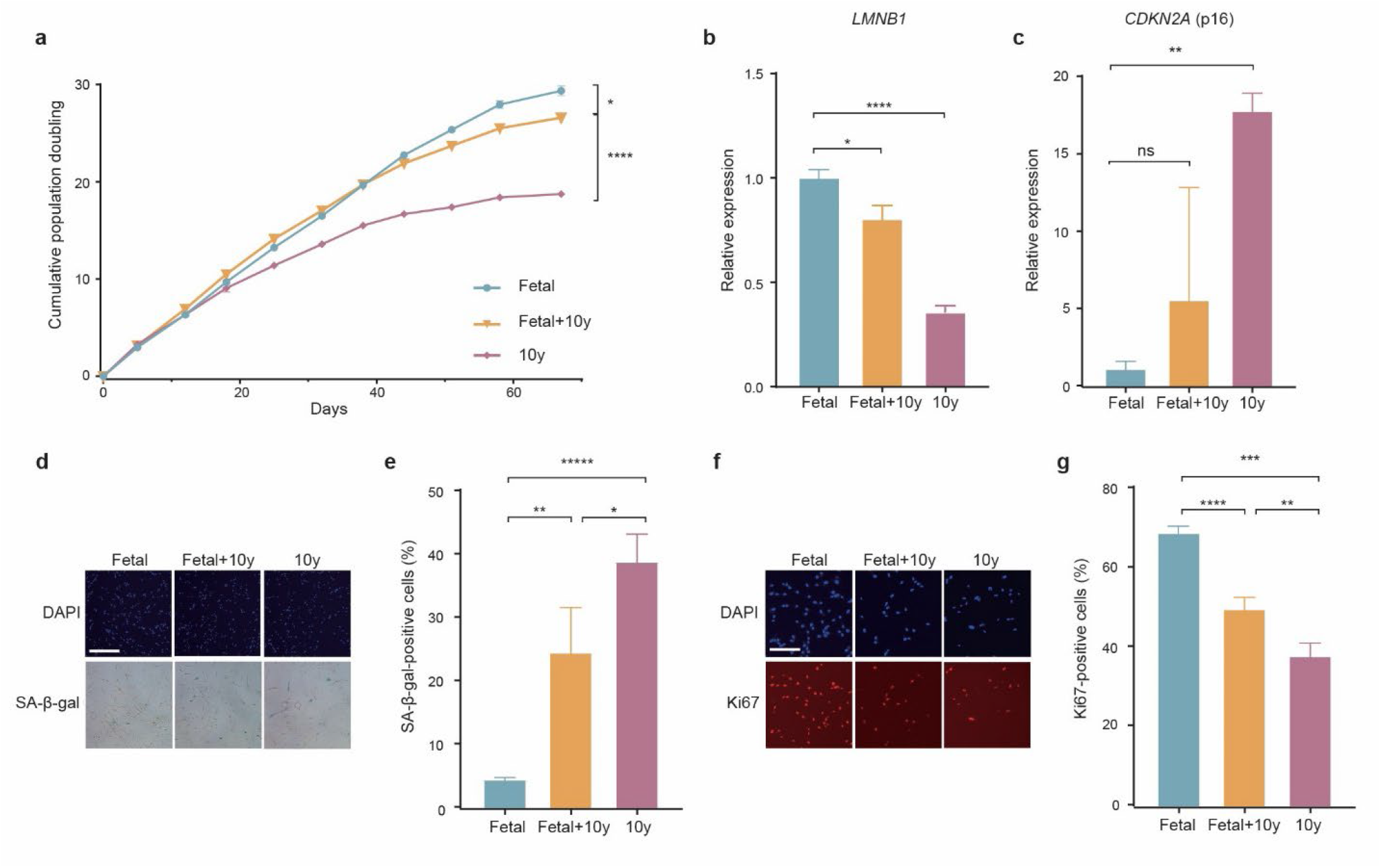
**a,** Replicative lifespan curves of IMR90 cultured in fetal (FBS), aged (10y), and heterochronic (50% fetal/50% aged) bovine serum. **b**, qPCR analysis of *LMNB1* in mid-late passage IMR90 cultured under different serum conditions, normalized to fetal serum condition. **c**, qPCR analysis of *CDKN2A* in mid-late passage IMR90 cultured under different serum conditions, normalized to fetal serum condition. **d-e**, Representative images (**d**) and quantification (**e**) of senescence-associated β-galactosidase (SA-β-Gal) staining of IMR90 at mid-late passage from different serum age conditions. **f-g,** Representative images (**f**) and quantification (**g**) of Ki67 staining of IMR90 at mid-late passage from different serum age conditions. Statistical analysis was performed using two-tailed unpaired *t*-tests (**a-c**,**e** and **g**). Data represented as mean ±SEM from three technical replicates, \**p*<0.05, \*\**p*<0.01, \*\*\**p*<0.001, \*\*\*\**p*<0.0001, ns=non-significant. Scale bars, 500 µm (**d**) and 200 µm (**f**).

**Extended Data Figure 2.**
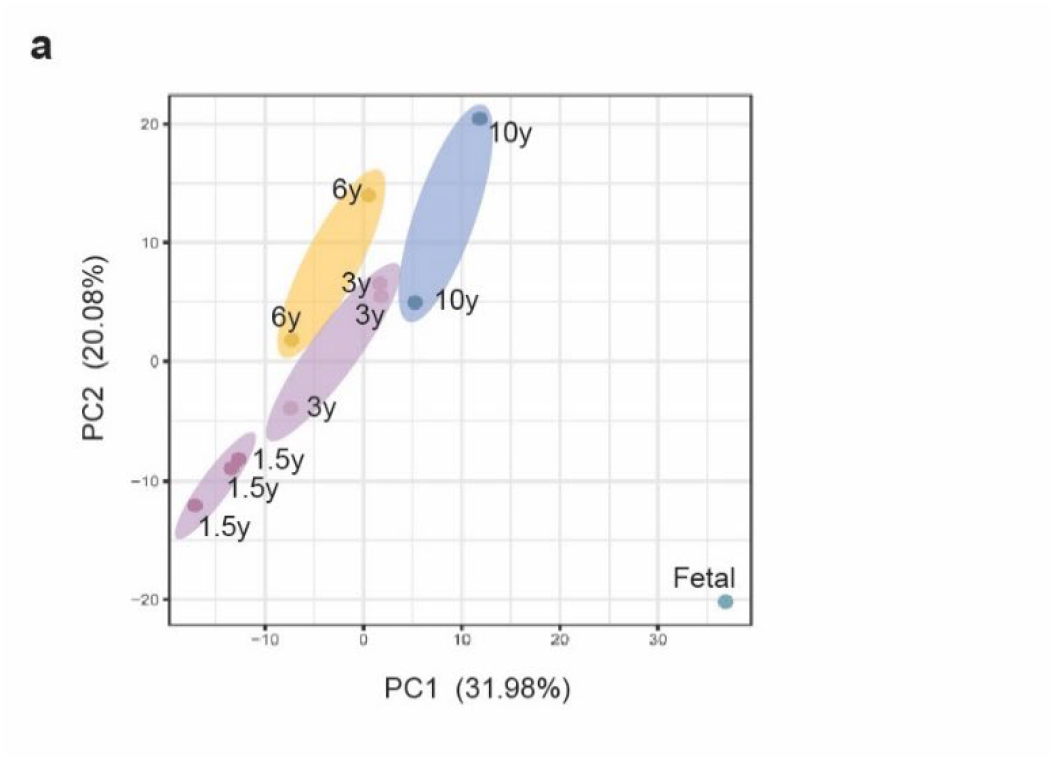
**a,** Principal component analysis (PCA) of protein signatures of different aged bovine serum.

**Extended Data Figure 3.**
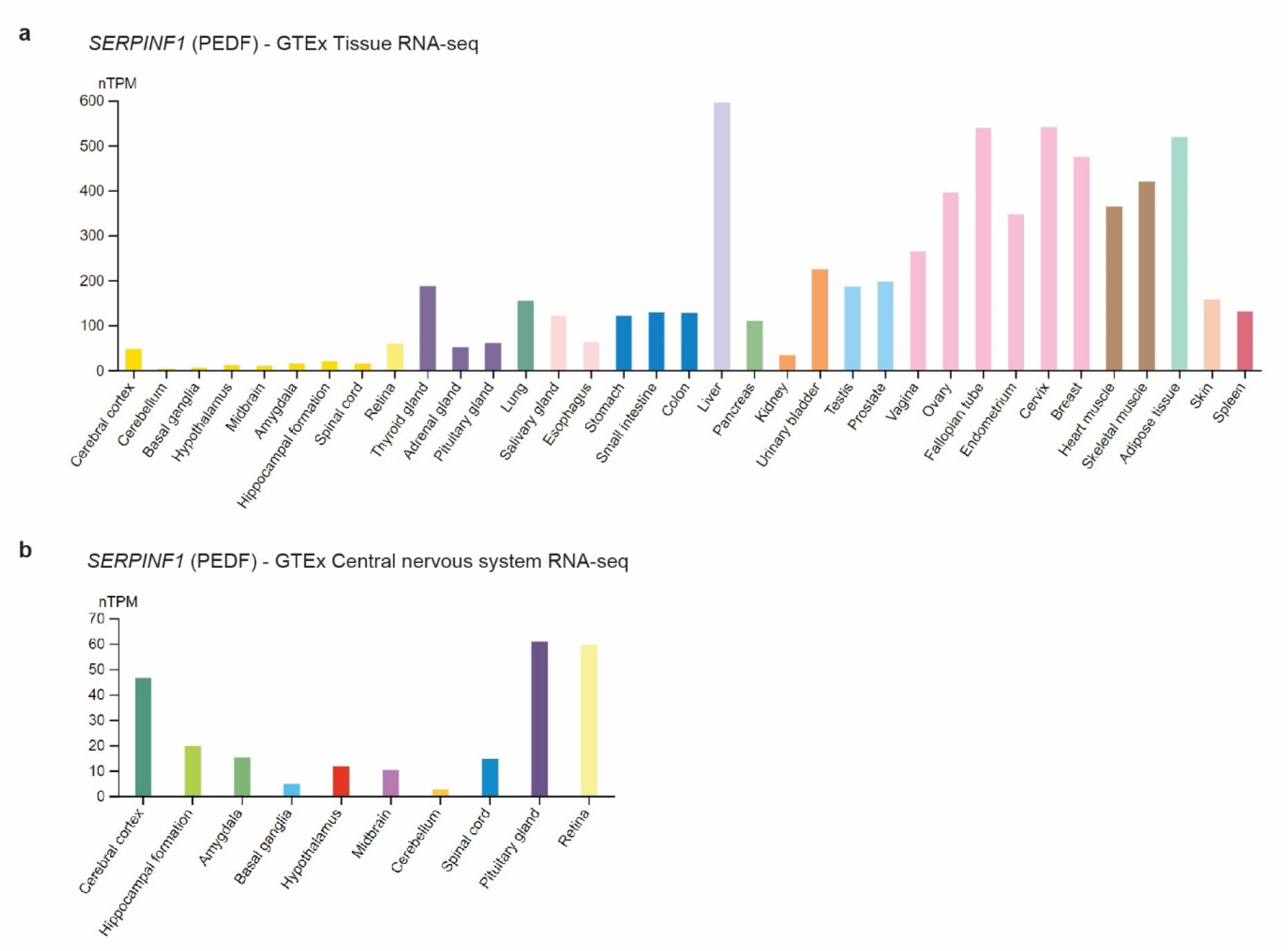
**a,** Expression of *SERPINF1* (PEDF) across human tissue from GTEx RNA-seq analysis. **b,** Expression of *SERPINF1* (PEDF) across human central nervous system tissue from GTEx RNA-seq analysis.

**Extended Data Figure 4.**
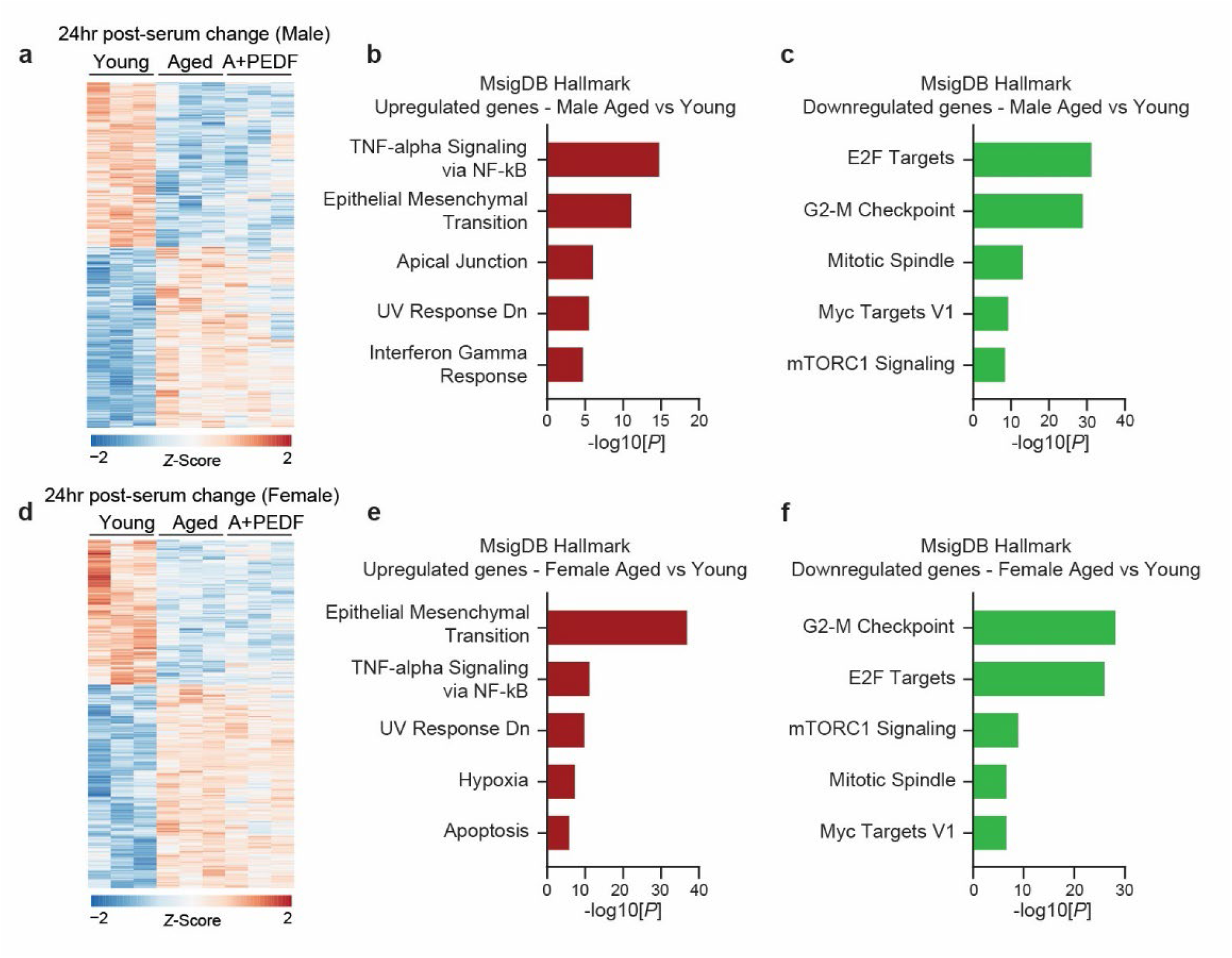
**a-c**, Significant DEGs (*p*<0.05) after RNA-seq analysis of IMR90 24hr post-serum change into either male young or aged serum, or aged serum supplemented with PEDF (**a**), and associated pathways (**b-c**). **d-f**, Significant DEGs (*p*<0.05) after RNA-seq analysis of IMR90 24hr post-serum change into either female young or aged serum, or aged serum supplemented with PEDF (**d**), and associated pathways (**e-f**). Statistical analysis was performed using Wald tests (**a** and **d**), and Fisher’s exact test (**b-c**, and **e-f**).

**Extended Data Figure 5.**
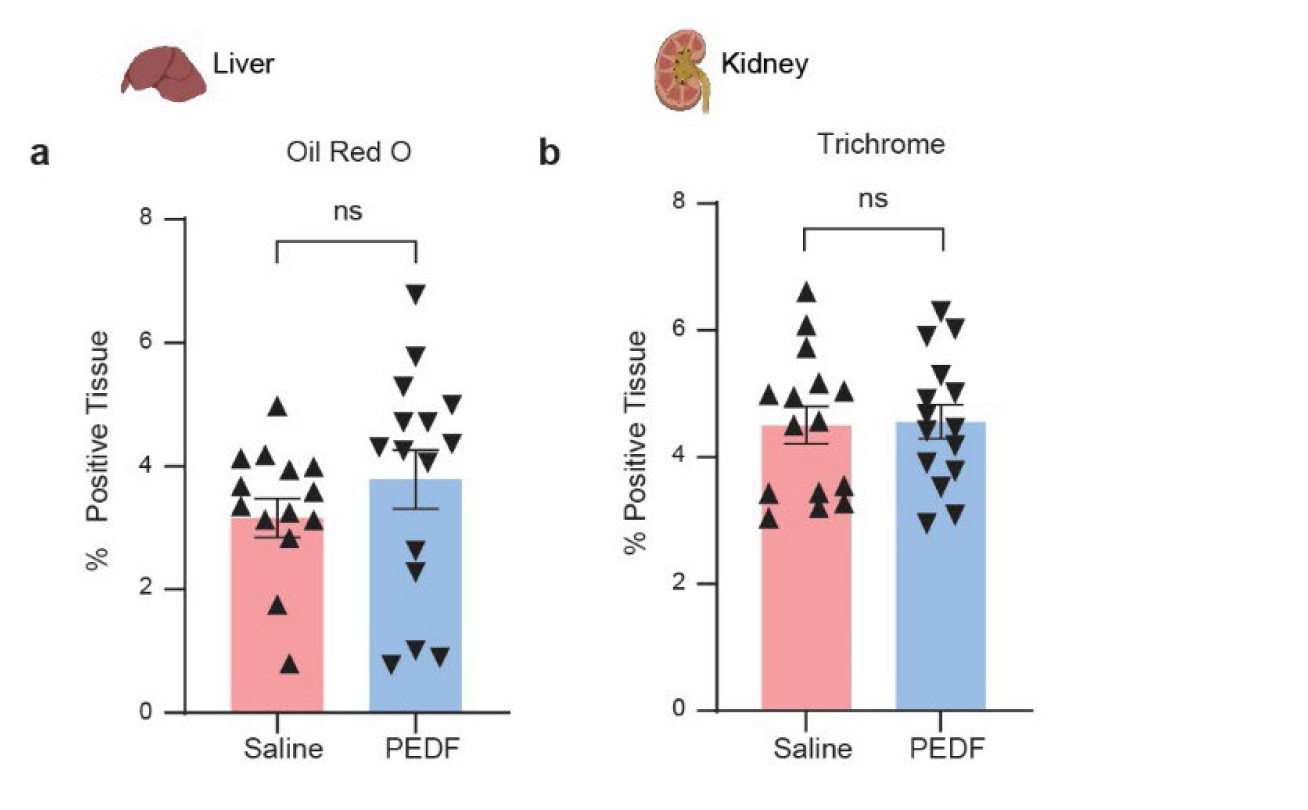
**a,** Quantification of Oil Red O staining in liver of aged mice (n=14 (saline), n=15 (PEDF) mice). **b,** Quantification of Trichrome staining in kidney of aged mice (n=15 (saline), n=15 (PEDF) mice). Statistical analysis was performed using two-tailed unpaired *t*-tests. Data represented as mean ±SEM, ns=non-significant.

## Methods

### IMR90 maintenance and subculturing for replicative life span assay

IMR90 cells were obtained from the American Type Culture Collection (ATCC, Lot# 64155514) at population doubling (PD) 25. The cells were grown in Eagle’s Minimum Essential Medium (MEM, Corning, 10-009-CV) supplemented with 10% fetal bovine serum (FBS, Gemini, 100-500, Lot A23G00J) and maintained in an incubator set to 37°C, 5% CO2, and 20% O2. 1×10^-6^ IMR90 were seeded and maintained on 10cm plates (Corning, 430293). Cells were split at 70% to 80% confluence. For IMR90 cells PD 25 to PD 40, it took 3 to 4 days to reach 80% confluence. Confluence levels were determined with a phase contrast EVOS XL Core microscope (Invitrogen). Late passage IMR90 cells were split every 5 to 7 days because of slower growth. Full media was replenished every other day. Cells were passaged by one time PBS wash, followed by dissociation with 0.05% Trypsin-EDTA (Gibco, 25300-54) for two to three minutes. Two times volume of full media was added to neutralize Trypsin. Cell suspension was collected into 15mL conical tube, followed by centrifugation at 300xg for 5 minutes. The supernatant was then aspirated. The cell pellet was resuspended in media and counted with C-Chip disposable hemocytometer (Incyto, DHC-N01-5). Cell numbers taken from the hemocytometer were used to calculate PDs. The following formula was used to calculate PDs: PD=log2(number of cells harvested/number of cells seeded). For the replicative life span assay, IMR90 PD 34 to PD 40 were seeded on 6 well plates (ThermoFisher, 140675) as the beginning points for the assays. The starting PDs within the same set of assays were identical. Cells for the same condition were tested in triplicates. All groups of cells within the same set of assays were maintained as described above, and passaged on the same day to avoid batch effects.

For the analyses of cell proliferation and cellular senescence markers, mid-late passage (PD38-59, depending on the culture conditions) cells were used. For RT-PCR analyses, additional plates were seeded during passaging. Cell pellets were subsequentially harvested 3 to 5 days later depending upon early passages or late passages.

### Serum

Serum samples used for all experimental procedures are listed in **Supplementary Table 9**.

### SA-β-gal staining

Cells were stained for SA-β-gal activity according to a previously described method ^87^.

### Immunofluorescent staining

For immunofluorescent staining cells were fixed with formaldehyde (4% in PBS) for 10 min, permeabilized with Triton X-100 (0.4% in PBS) for 15 min, incubated with blocking buffer (10% donkey serum in PBS) for 1 hour, and stained with fluorescent APC conjugated anti-Human Ki67 antibody (eBioscience, 17-5699-42), with 1% donkey serum overnight at 4 °C. The cells were washed three times with PBS. Hoechst 33342 (Invitrogen) was used to stain nuclear DNA.

### RT-qPCR

Total RNA was extracted from cells using RNeasy Plus Mini Kit (Qiagen, 74136). 0.5-1.0 μg total RNA was used for cDNA synthesis with 5x PrimeScript RT Master Mix (Takara, RR036A-1). Quantitative real-time PCR was carried out using PowerUp SYBR Green Master Mix (Applied Biosystems, 100029284) on a QuantStudio 6 Pro (Applied Biosystems). All data were normalized by 18S rRNA transcript and calculated using the ΔΔCq method. All RT-qPCR primer pairs are listed in **Supplementary Table 10**.

### Tissue collection

Animals were anesthetized with isoflurane and transcardially perfused with PBS and tissues were harvested and split into samples. Samples processed for histology were fixed overnight at 4 °C in 4% paraformaldehyde in PBS (4%PFA). Fixed tissues processed for cryopreservation were washed with PBS and incubated with 15% sucrose in PBS overnight at 4 °C then 30% sucrose in PBS overnight at 4 °C. Sucrose treated tissue was embedded in optimum cutting temperature in cryomolds (Tissuetek) and frozen in 2-methylbutane prechilled in liquid nitrogen. Frozen embedded tissue was cryosectioned at 10 μm thickness, collected on Superfrost Plus microslides (Fisherbrand) and stored at 80 °C until analysis. For RNA-seq analysis the hippocampus was subdissected, whole liver and kidney was collected, and tissues were snap-frozen.

### RNA isolation and bulk RNA-seq

Total RNA was extracted from fresh frozen cells and tissue samples using Qiagen RNeasy Plus Universal mini kit following manufacturer’s instructions (Qiagen). RNA samples were quantified using Qubit 2.0 Fluorometer (Life Technologies) and RNA integrity was measured using the RNA Screen Tape on Agilent 2200 TapeStation (Agilent Technologies). Samples were initially treated with TURBO DNase (Thermo Fisher Scientific) to remove DNA contaminants. The next steps included performing rRNA depletion using QIAseq FastSelect−rRNA HMR kit (Qiagen), which was conducted following the manufacturer’s protocol. RNA sequencing libraries were constructed with the NEBNext Ultra II RNA Library Preparation Kit for Illumina by following the manufacturer’s recommendations. Briefly, enriched RNAs are fragmented for 15 minutes at 94 °C. First strand and second strand cDNA are subsequently synthesized. cDNA fragments are end repaired and adenylated at 3’ends, and universal adapters are ligated to cDNA fragments, followed by index addition and library enrichment with limited cycle PCR. Sequencing libraries were validated using the Agilent Tapestation 4200 (Agilent Technologies) and quantified using Qubit 2.0 Fluorometer (Thermo Fisher Scientific) as well as by quantitative PCR (KAPA Biosystems). The sequencing libraries were multiplexed and clustered onto a flowcell on the Illumina NovaSeq instrument according to manufacturer’s instructions. The samples were sequenced using a 2×150bp Paired End (PE) configuration. Image analysis and base calling were conducted by the NovaSeq Control Software (NCS). Raw sequence data (.bcl files) generated from Illumina NovaSeq was converted into fastq files and de-multiplexed using Illumina bcl2fastq 2.20 software. One mis-match was allowed for index sequence identification. Transcript expression was quantified from RNA-seq reads using salmon (v.1.10) and either the human Gencode v43 or mouse Gencode vM34 gene models. Differential expression analysis was performed in R (v.4.3.2) by Wald test in the DESeq2 package^88^ (v.1.42.0). Genes significantly changed *in vitro* from exposure to different culture conditions were determined using an FDR < 0.05, and significance in mouse tissue after systemic PEDF administration was determined with a nominal p < 0.05. Hallmark gene sets (MSigDB Hallmark 2020) and Gene Ontology (GO Biological Process 2023) enrichment analysis was performed using Enrichr^89^.

### Mass spectrometry

Label free data-independent acquisition (DIA) quantitative proteomics was performed on bovine serum by BGI (Cambridge, MA). Serum sample preparation: 6 μL (about 300 μg) of each sample was taken for digestion by adding 25% sodium deoxycholate to 4% final concentration and dithiothreitol (DTT) to 10 mM final concentration, incubated at 60 °C for 30 minutes, followed by addition of iodoacetamide (IAM) to 30 mM and incubation at room temperature for 30 minutes. Additional DTT was added to bring to 30mM to quench excess iodoacetamide. 50 mM ammonium bicarbonate was added to bring sodium deoxycholate concentration to 1% and 8 μg of LysC/Trypsin mixture (Thermo Fisher Scientific) was added to each sample for overnight digestion at 37 °C. After digestion, samples were acidified with trifluoroacetic acid to 0.4%, precipitating the sodium deoxycholate. The samples were spun at 15,000 x g for 5 minutes and the resulting supernatants were taken for cleanup. Peptide Cleanup: 10% TFA was added into each sample of digested peptides samples and formed a final concentration of 1% TFA. pH was tested and acidic. Then, acidified samples went through 100 mg SEK PAK columns (Cat No: 60108-302, ThermoFisher Scientific) for desalting. About 1/3 of each sample was taken to pool together for peptide fractionation; remaining 2/3 of each sample was dried and stored for LC-MS/MS analysis. Peptide Fractionation: The DIA library composite samples were fractionated into 96 fractions with a high pH reverse phase offline HPLC fractionator (VanquishTM, ThermoFisher Scientific). Mobile phase A is DI H2O with 20 mM Formic Acetate, pH 9.3; mobile phase B is Acetonitrile (OptimaTM, LC/MS grade, Fisher ChemicalTM) with 20mM Formic Acetate, pH 9.3. 96 fractions were then combined into 24 fractions and were ready for Liquid Chromatography Mass Spectrometry (LC/MS) analysis.

### LC-MS/MS Analysis

All fractionated samples were analyzed by nano flow HPLC (Ultimate 3000, Thermo Fisher Scientific) followed by Thermo Orbitrap Mass Spectrometer (QE HF-X). Nanospray FlexTM Ion Source (Thermo Fisher Scientific) was equipped with Column Oven (PRSO-V2, Sonation) to heat up the nano column (PicoFrit, 100 µm x 250 mm x 15 µm tip, New Objective) for peptide separation. The nano LC method was water acetonitrile based and 150 minutes long with a 0.25 µL/min flowrate. For each library of fractions, all peptides were first engaged on a trap column (Cat. No: 160454, Thermo Fisher) and then were delivered to the separation nano column by the mobile phase. For DDA library construction, a DIA library specific DDA MS2-based mass spectrometry method on Eclipse was used to sequence fractionated peptides that were eluted from the nano column. For the full MS, 120,000 resolution was used with the scan range of 375 m/z – 1500 m/z. For the dd-MS(MS2), 15,000 resolution was used, and Isolation window was 1.6 Da. ‘Standard’ AGC target and ‘Auto’ Max Ion Injection Time (Max IT) were selected for both MS1 and MS2 acquisition. Collision Energy (NCE) was set to 35%, and total cycle time was 1 sec. For DIA analytical samples, a high-resolution full MS scan followed by two segment DIA methods was used for the DIA data acquisition. For the full MS scan, 120,000 resolution was used for the range of 400 m/z – 1200 m/z with ‘Standard’ AGC target and 50 ms Max IT. For DIA fragments scan, 30,000 resolution was used for the range of 110 m/z – 1,800 m/z with ‘Standard’ AGC target and ‘Auto’ Max IT.

### Generation of recombinant PEDF

The human PEDF gene (*SERPINF1*) was cloned in pcDNA3.1+C-6His by XhoI / ApaI and confirmed by Sanger sequencing. HEK-293 cells were transfected with the expression vector and secreted protein was purified by affinity chromatography (His60 resin, Clontech) and size exclusion chromatography (26/60 Superdex 200 HiLoad (GE), PBS mobile phase), tested for endotoxin and analyzed by Coomassie stained SDS-PAGE. Protein identity was confirmed by mass spectrometry.

### Animal models

Aged male C57BL/6 mice were obtained from the National Institute on Aging aged rodent colony and were randomly assigned to experimental groups. All mice were kept on a 12-h light/dark cycle and provided ad libitum access to food and water. Mice were handled according to the National Institutes of Health’s Guide for the Care and Use of Laboratory Animals. All procedures involving mice were approved by the Columbia University Institutional Animal Care and Use Committee. Recombinant PEDF was dosed at 4 µg day ^-1^ using subcutaneously implanted Alzet mini-osmotic pumps (model 2006). Control animals were implanted with Alzet pumps containing saline (PBS; Corning).

### Novel object recognition

Novel object recognition measures an animals’ ability to differentiate a novel object from a familiar object to test explicit memory. Tests were conducted in a clean white rectangular arena. On Day 1, the animal was habituated to the arena for 30 minutes. On Day 2 (testing day), the animal was habituated to the arena again, for 30 minutes. The subject was then briefly removed from the arena while the experimenter placed two identical objects in the arena. The subject was then returned to the arena and allowed to freely explore the two objects for 10 minutes (for familiarization). An hour after the familiarization session, the subject was returned to the arena for a 5-min memory test. For the memory test, one of the objects used in the familiarization session was replaced with a novel object. Time spent interacting with each object was analyzed from recorded videos, by investigators blind to treatment information. Object investigation was defined as time spent sniffing the object when the nose is oriented toward the object and the nose-object distance is 2 cm or less. Novel object memory was defined as spending significantly more time sniffing the novel object than the familiar object. Failing to show preference for the novel object was interpreted as a memory deficit. The toys were washed with soap and hot water and dried with paper towel after each use. The arena was cleaned with 70% ethanol and wiped dry with paper towels after each session. Novelty preference was determined by calculating preference=time spent with novel object/(time spent with novel object + time spent with familiar object)*100.

### Mouse tissue histology

Tissue samples were prepared in OCT in cryomolds. Slides were stained by standard PicroSirius Red and Oil Red O protocols (Histowiz, Brooklyn NY). Sections were counterstained with hematoxylin, dehydrated, and film-coverslipped (Sakura TissueTek-Prisma Coverslipper). Whole slide scanning (40x) was performed using a Leica Aperio AT2 slide scanner (Leica Microsystems) at 40X.

### Statistical and bioinformatic analyses

Statistical analyses of differences in replicative lifespan and molecular and cellular markers of senescence were via two-tailed unpaired *t*-test. To analyze the changes in protein levels throughout the aging process, the plasma protein levels were standardized using z-scores.

Locally estimated scatterplot smoothing (LOESS) regression was then applied to each plasma factor to estimate their trajectories. To identify groups of proteins with similar trajectory patterns, pairwise differences between LOESS estimates were computed using Euclidean distance.

Hierarchical clustering was performed using the complete method to cluster the proteins based on their trajectory patterns. To detect the effect of age on protein abundance a linear model was fitted in R, using the function “lm(age ∼ protein expression)”. Statistical significance was subjected to multiple testing correction by using “p.adjust(p-value, method=”BH”)” function in R with an FDR cutoff of p<0.05.

## Data availability

DIA mass spectrometry raw data will be deposited in the ProteomeXchange Consortium. All bulk RNA-seq data will be available at the Gene Expression Omnibus (GEO).

## Notes

### Competing Interest Statement

The authors have declared no competing interest.

